# Deterministic versus Probabilistic Tractography: Impact on White Matter Bundle Shape

**DOI:** 10.1101/2025.08.19.669949

**Authors:** Yuhan Shuai, Bramsh Qamar Chandio, Yixue Feng, Julio E. Villalon-Reina, Talia M. Nir, Christopher R. K. Ching, Iyad Ba Gari, Sophia I. Thomopoulos, Jonathan Davis Alibrando, John P. John, Ganesan Venkatasubramanian, Neda Jahanshad, Paul M. Thompson

## Abstract

In diffusion MRI-based tractography, deterministic and probabilistic algorithms reconstruct white matter using distinct strategies, yet their impact on bundle morphology remains uncertain. Using bundle shape similarity analysis, we compared both methods for the left arcuate fasciculus (AF_L) (The left arcuate fasciculus is a critical white matter tract that connects language comprehension and production areas in the human brain, enabling fluent language processing) across four datasets: Alzheimer’s Disease Neuroimaging Initiative (ADNI), Human Connectome Project-Aging (HCP-A), National Institute of Mental Health and Neurosciences (NIMHANS), and Pediatric Imaging, Neurocognition, and Genetics (PING). Probabilistic tractography consistently produced higher inter-subject shape similarity, by capturing broader anatomical trajectories and enhancing reproducibility. However, this extensive coverage may obscure subtle pathological variations critical for clinical detection. Bundle shape similarity analysis with atlas corroborated these findings, showing stronger alignment for probabilistic tracking and highlighting its utility in quantitative quality control. These results emphasize the need to balance morphological consistency with sensitivity to neuroanatomical variation when selecting tractography methods for research and clinical applications.

## I. Introduction

Diffusion-weighted magnetic resonance imaging (dMRI) provides a noninvasive method to probe white matter organization in the human brain by capturing the directional dependence of water diffusion along fiber tracts. Building on this principle, dMRI-based fiber tractography reconstructs long-range pathways from local fiber orientations, enabling detailed mapping of white matter architecture for studying brain anatomy, development, and clinical applications [1]. However, there remains debate regarding the optimal methods for reconstructing fiber pathways. In particular, the choice between deterministic [2], [3] and probabilistic [4], [5] tractography algorithms significantly influences which streamlines are generated and how far they propagate.

Deterministic tractography relies on voxel-wise diffusion model estimates and propagates streamlines along the principal diffusion direction. As a result, streamlines originating at a given seed point will always follow the same trajectory, producing consistent paths that represent the dominant orientation [2], [3]. In contrast, probabilistic tractography explicitly accounts for uncertainty in local fiber orientations by sampling from a distribution of possible directions at each step. This approach allows for greater exploration of alternative pathways but introduces variability, as repeated tracking from the same seed may produce different trajectories. By modeling these uncertainties, probabilistic tractography can capture a wider range of possible anatomical configurations [4], [5]. A critical question, however, is how these algorithmic differences affect the morphological consistency of reconstructed white matter bundles across individuals. To address this, we adopted a bundle shape-based evaluation framework, quantifying similarity using the bundle adjacency (BA) metric, which ranges from 0 (no shared shape features) to 1 (identical shape). BA provides a robust measure of 3D morphological similarity, enabling us to assess whether deterministic tractography results more uniform bundle shapes compared to the potentially more variable probabilistic reconstructions. In this study, we compared deterministic and probabilistic tractography in terms of bundle shape similarity across four heterogeneous datasets: ADNI (n=10), HCP-A (n=5), PING (n=5), and NIMHANS (n=10). These cohorts span diverse age groups, imaging protocols, and clinical contexts (e.g., ADNI includes older adults with varying cognitive status, whereas PING represents a younger population), providing a comprehensive testbed for generalizability. We generated whole-brain tractograms for each subject using local deterministic and probabilistic algorithms with identical parameters and extracted major white matter tracts. We then computed inter-subject shape similarity for each bundle within each cohort, focusing on the left arcuate fasciculus (AF_L) as a representative tract due to its anatomical complexity and variability across individuals, making it a sensitive benchmark for evaluating tractography performance [6]. We hypothesize that deterministic tractography produces more shape-consistent bundles across individuals due to its reliance on principal diffusion directions. In contrast, probabilistic tractography introduces greater morphological variability through uncertainty sampling and exploration of alternative pathways.

## II. Methods

We analyzed diffusion MRI data from four cohorts with varying demographics and imaging parameters: (1) ADNI – 10 older adults from the Alzheimer’s Disease Neuroimaging Initiative (age ∼ 50–85, cognitively normal/mild cognitive impairment), (2) HCP-A – 5 adults from the Human Connectome Project Aging cohort (age ∼ 50–75, healthy aging), (3) PING – 5 children and adolescents from the Pediatric Imaging, Neurocognition, and Genetics cohort (age ∼ 3–20, healthy developing brains), (4) NIMHANS – 10 adults from a NIMHANS research dataset (age ∼ 50–85, dementia/mild cognitive decline/cognitive normal). All diffusion datasets underwent standard preprocessing, including motion and eddy-current correction, brain masking, and diffusion tensor estimation or fiber orientation reconstruction as appropriate.

### A. Tractography Algorithm

Whole-brain tractograms were generated twice for each subject using local deterministic and probabilistic tracking in DIPY [7], with tracking parameters held constant to isolate algorithmic effects. Fiber orientation distributions were estimated using the single-shell single-tissue (SSST) or multi-shell multi-tissue constrained spherical deconvolution (CSD) model [8], depending on the dataset. Tracking was seeded from voxels with FA *>* 0.15 (8 seeds per voxel), propagated with a 0.5 mm step size, and terminated when FA *<* 0.15. All seeding strategies, stopping criteria, and anatomical constraints were identical across reconstructions, differing only in the tracking algorithm (deterministic vs. probabilistic).

### B. Bundle Extraction and Shape Similarity Analysis

Whole-brain tractograms were affinely registered to an Atlas in MNI space using streamline-based registration (SLR) method [9]. White matter tracts were extracted from whole-brain tractograms using an auto-calibrated version of RecoBundles [10], [11], which used HCP1065 atlas bundles as a reference for segmenting streamlines from the tractograms. We applied auto-calibrated RecoBundles with identical parameters for both deterministic and probabilistic tractograms, ensuring that the bundle segmentation process itself did not bias one method over the other.

We then quantified bundle shape similarity across bundles using the network-based bundle shape similarity approach proposed in BUndle ANalytics (BUAN) framework [11]. The bundle shape similarity method uses bundle adjacency (BA) metric [13] to quantify the resemblance between two bundles based on the minimum direct flip (MDF) [13] distance between their constituent streamlines. The MDF distance is a symmetric measure that accounts for streamline bi-directionality by computing both direct and flipped distances and taking the minimum of the two. The BA metric defines adjacency between two streamlines *b*_1_ ∈ *B*_1_ and *b*_2_ ∈ *B*_2_ if their MDF distance satisfies *MDF* (*b*_1_, *b*_2_) ≤ *θ*, with *θ >* 0 being a predefined threshold. The coverage of *B*_1_ by *B*_2_ is then computed as the fraction of streamlines in *B*_1_ that are adjacent to at least one streamline in *B*_2_. Similarly, coverage of *B*_2_ by *B*_1_ is calculated. The symmetric BA measure is defined as the average of the two coverage values:

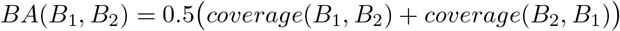

These scores range from 0 (no shared shape characteristics) to 1 (identical shape). In our analyses, we set strict similarity threshold of 5 mm. First, within each dataset, fully connected shape similarity network was computed by calculating pair-wise between all subjects for both deterministic and probabilistic tractography methods, resulting in a symmetric similarity matrix for tracts of each approach (e.g., 10 subjects yielding a 10 × 10 matrix). Heatmaps were generated to visualize these matrices, enabling qualitative assessment of intra-cohort shape variability and quantitative comparison between deterministic and probabilistic tracking outcomes. After computing within-cohort shape similarity, each subject’s bundle was compared to its corresponding standardized HCP1065 atlas bundle (used in bundle segmentation as reference model bundle). For statistical evaluation, paired t-tests were conducted to compare mean within-cohort BA scores between deterministic and probabilistic tractography. Statistical significance was assessed at an *α* level of 0.05.

## III. Results

### A. Inter-subject Bundle Shape Similarity Analysis

Comparison across tractography methods revealed notable variability in inter-subject shape similarity for the AF_L bundle (see Table 1). Across most datasets, probabilistic tracking produced higher inter-subject BA compared with deterministic tracking. In the ADNI cohort (*n* = 10), mean BA increased from 0.56 ± 0.09 (deterministic) to 0.78 ± 0.05 (probabilistic), a highly significant difference (paired *t*-test: *t*(9) = − 6.40, *p <* 0.001; Wilcoxon signed-rank: *W* = 0, *p* = 0.002; Cohen’s *d* = 2.02). A similar trend was observed in HCP-A (*n* = 5), where BA rose from 0.77 ± 0.05 to 0.82 ± 0.06 (*t*(4) = − 4.63, *p* = 0.0098; Wilcoxon *p* = 0.063; *d* = 2.07), and in NIMHANS (*n* = 10), where the effect was even larger (0.58 ± 0.08 vs. 0.83 ± 0.04; *t*(9) = − 9.72, *p <* 0.00001; Wilcoxon *p* = 0.002; *d* = 3.07). In contrast, the PING dataset did not show significant differences between the methods (0.65 ± 0.07 vs. 0.63 ±0.14; *t*(4) = 0.59, *p* = 0.588; Wilcoxon *p* = 1.0).

**TABLE I:** Inter-Subject BA for AF_L Across Datasets.

| Dataset | Method | Mean BA ± SD | BA Range |
| --- | --- | --- | --- |
| ADNI | Deterministic | $0.56 \pm 0.09$ | 0.43–0.68 |
| | Probabilistic | $0.78 \pm 0.05$ | 0.69–0.86 |
| HCP-A | Deterministic | $0.77 \pm 0.05$ | 0.71–0.85 |
| | Probabilistic | $0.82 \pm 0.06$ | 0.71–0.94 |
| NIMHANS | Deterministic | $0.58 \pm 0.08$ | 0.48–0.77 |
| | Probabilistic | $0.83 \pm 0.04$ | 0.73–0.89 |
| PING | Deterministic | $0.65 \pm 0.07$ | 0.54–0.75 |
| | Probabilistic | $0.63 \pm 0.14$ | 0.40–0.86 |

### B. Visualization of Bundle Similarity within Dataset

To illustrate method-specific patterns of inter-subject similarity, Figure 3 presents pairwise similarity matrices for the AF_L in the ADNI cohort as an example. The deterministic matrix (left) exhibits pronounced variability across subject pairs, with several lighter regions indicating lower similarity. By contrast, the probabilistic matrix (right) shows more uniform and darker tones, reflecting higher and more consistent similarity scores.

**Fig. 1:**
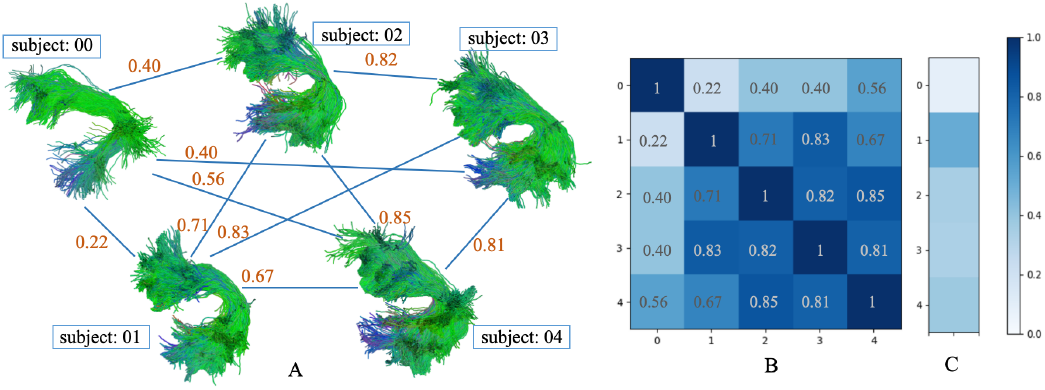
Fully connected bundle shape similarty network (A), adjacency matrix (B) and comparison with HCP1065 atlas (C) of the AF_L bundle in PING dataset (probabilistic tracking).

**Fig. 2:**
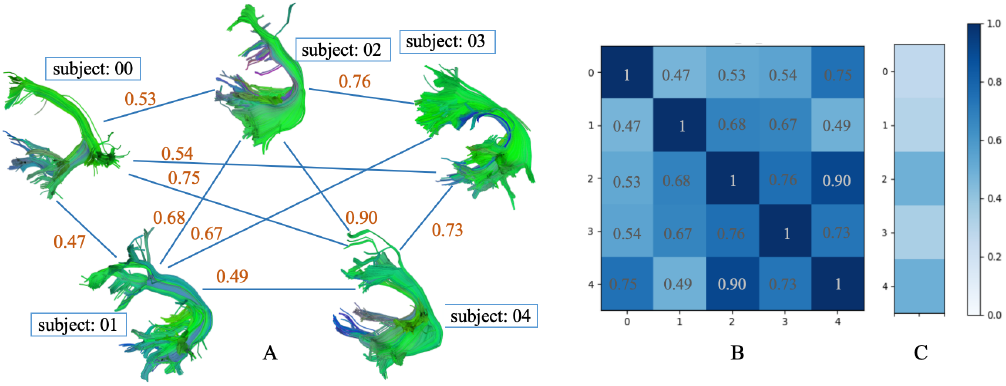
Fully connected bundle shape similarty network (A), adjacency matrix (B) and comparison with HCP1065 atlas (C) of the AF_L bundle in PING dataset (deterministic tracking).

**Fig. 3:**
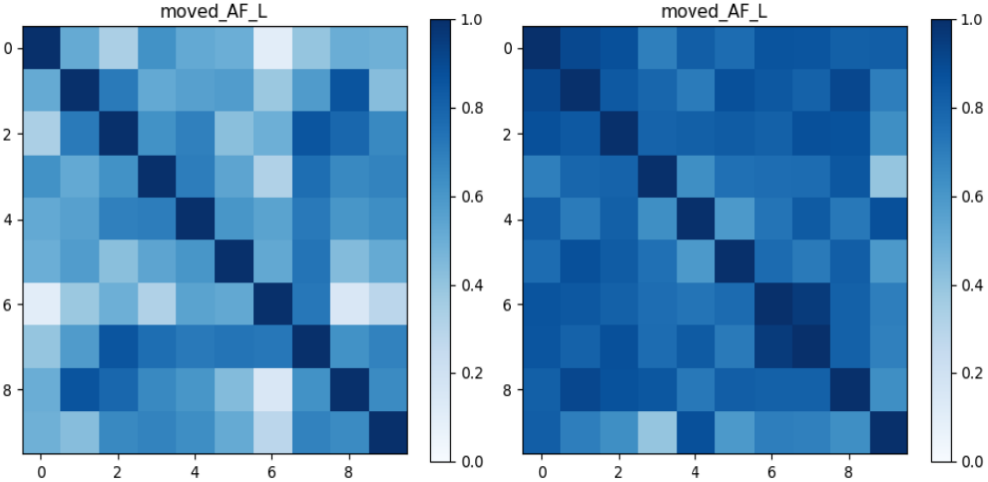
Shape similarity matrices for the AF_L bundle in the ADNI dataset using deterministic (left) and probabilistic (right). Each matrix shows pairwise shape similarity scores across subjects, with darker colors indicating higher similarity.

Figure 4 provides 3D anatomical visualization of AF_L tractography in a representative ADNI subject under both tracking algorithms, compared with the HCP1065 reference atlas. Probabilistic tracking (middle) captures more continuous and widespread fibers relative to deterministic tracking (left), which appears sparser and less coherent. This difference highlights how algorithmic choices influence both quantitative similarity metrics and qualitative tract geometry.

**Fig. 4:**
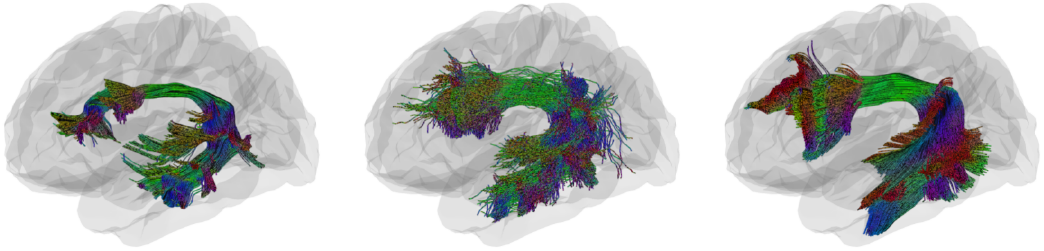
Example of tractometry visualization of the AF_L in a representative ADNI subject compared to the HCP-based atlas. Panels show (left) deterministic tracking, (middle) probabilistic tracking, and (right) reference image from HCP atlas.

### C. Bundle Similarity Comparing with HCP1065 atlas

Atlas-based BA values for the AF_L are summarized in Table II. In the ADNI cohort, deterministic tracking produced a mean BA of 0.33 ± 0.14 (range 0.07–0.57), while probabilistic tracking produced 0.37 ± 0.08 (0.22–0.47). A paired comparison of subject-level BA scores revealed no significant difference between methods (*t*(9) = − 0.71, *p* = 0.497; Wilcoxon *p* = 0.322; Cohen’s *d* = − 0.22). For the HCP-A dataset, deterministic tracking achieved higher atlas alignment (0.67 ± 0.08) than probabilistic tracking (0.60 ± 0.13; *t*(4) = 2.97, *p* = 0.041; Wilcoxon *p* = 0.063; *d* = 1.33), indicating a modest effect favoring the deterministic approach. In contrast, NIMHANS exhibited slightly greater BA for probabilistic tracking (0.47 *±* 0.11) compared with deterministic (0.38 *±* 0.19), but this difference did not reach statistical significance (*t*(9) = −1.55, *p* = 0.158; Wilcoxon *p* = 0.203; *d* = −0.49). Lastly, PING showed comparable performance between methods (0.33 *±* 0.15 vs. 0.38 *±* 0.10; *t*(4) = −0.62, *p* = 0.565; Wilcoxon *p* = 0.625; *d* = −0.28).

**TABLE II:**
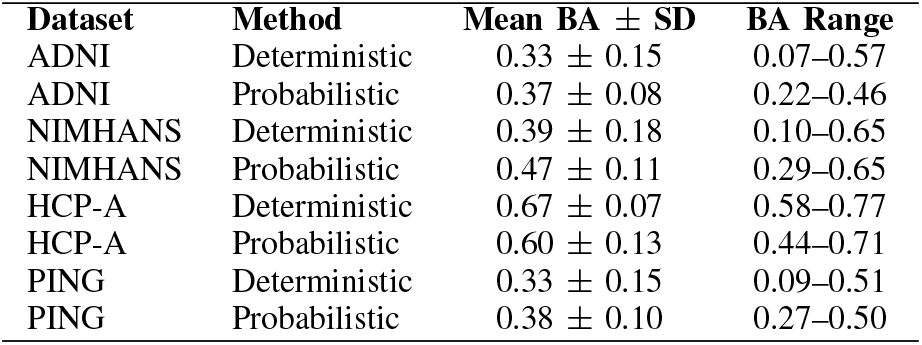
Atlas-based BA across Datasets.

Figure 5 illustrates atlas-based shape similarity for the AF_L bundle using representative subjects from the ADNI and HCP-A datasets, where darker colors indicate higher similarity with the atlas. The ADNI subject shows moderate correspondence, with probabilistic tracking exhibiting slightly more uniform coverage, whereas deterministic tracking displays localized regions of high similarity. In contrast, the HCP-A subject exhibits consistently darker patterns across both methods, reflecting stronger alignment with the HCP atlas.

**Fig. 5:**
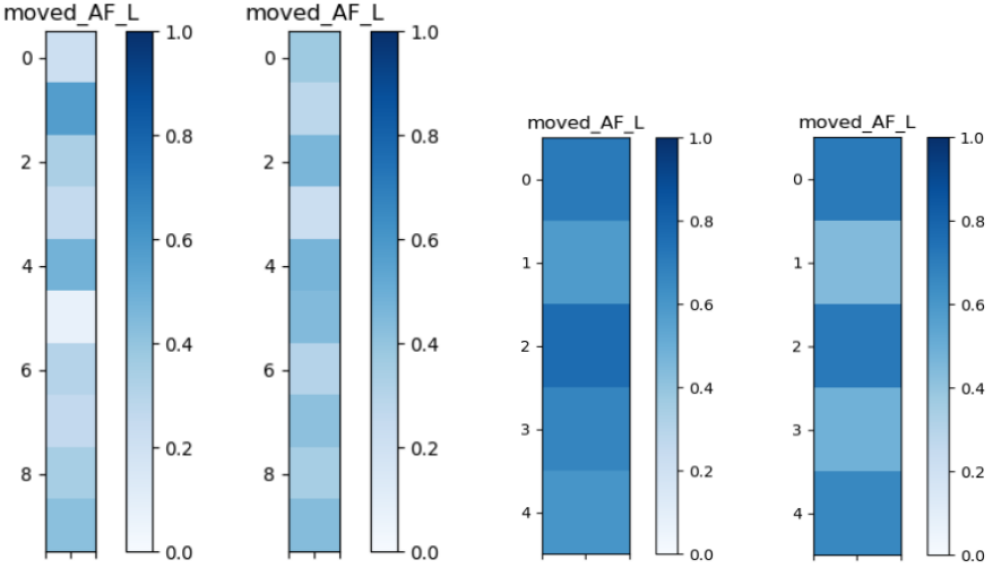
Example visualization of the AF_L in a representative ADNI subject compared to the HCP-based atlas. Each panel show (left) deterministic tracking, (right) probabilistic tracking.

## IV. Discussion and Conclusion

In this study, we compared deterministic and probabilistic tractography algorithms using a morphology-based approach, specifically focusing on the left arcuate fasciculus across diverse populations. Our findings consistently showed higher inter-subject BA scores for probabilistic tractography compared to deterministic tracking in ADNI, HCP-A, and NIMHANS datasets. This suggests that by incorporating directional uncertainty and sampling broader anatomical trajectories, probabilistic algorithms produce more consistent bundle reconstructions across individuals. Our results also indicate that probabilistic methods achieve greater morphological reproducibility by reliably capturing core and peripheral fibers. Deterministic methods, conversely, demonstrated lower consistency, likely owing to their susceptibility to local fiber orientation uncertainty and subsequent truncations.

These methodological differences exhibited notable age- and cohort-dependent patterns. Probabilistic tracking showed clear superiority in datasets comprising healthy adults (HCP-A), likely reflecting more stable and mature white matter structures. Conversely, in younger developmental populations (PING), deterministic and probabilistic methods resulted comparable shape consistency, potentially due to ongoing fiber maturation and inherently higher anatomical variability at younger ages [14]. Probabilistic methods again demonstrated enhanced consistency in the ADNI cohort, which includes cognitively normal individuals, mild cognitive impairment, and dementia patients. However, interpreting these results warrants caution. While probabilistic tracking effectively captures normal anatomical variability, its broader fiber inclusion might inadvertently obscure subtle pathological alterations, which often present as deviations from typical fiber trajectories [15]. Consequently, despite its lower overall consistency, deterministic tracking might preserve clinically relevant morphological variability indicative of early or subtle pathology, features potentially masked by probabilistic averaging [16], [17].

Atlas-based analyses further supported these conclusions, demonstrating that probabilistic tracking achieved better conformity with standardized anatomical references such as in the ADNI and NIMHANS cohorts. Incorporating quantitative atlas-based morphological comparisons thus offers an objective complement to visual quality control (QC), enhancing reproducibility and reliability of bundle reconstructions. Indeed, visual QC inherently evaluates morphological congruence between reconstructed bundles and known anatomical structures. Quantitative shape metrics, such as BA, could therefore augment existing QC protocols by automatically flagging reconstructions that deviate significantly from anatomical standards.

Our analysis focused exclusively on the AF_L bundle, which limits the generalizability of our findings to other tracts. Future work should expand morphological evaluations to additional bundles and larger, more diverse datasets. In parallel, we will perform along-tract BUAN tractometry [11] to map DTI-derived microstructural metrics such as FA and MD along white matter trajectories, enabling a more comprehensive assessment of how tractography algorithms influence tractometry outcomes. Integrating these microstructural indices will allow us to better characterize tractography performance across populations and improve the interpretability of downstream analyses. While parameters were held constant to isolate algorithmic effects, tractography results can still vary with factors like seeding, FA threshold, and step size, warranting sensitivity analyses to assess whether differences arise from algorithms or parameter choices.

In conclusion, our morphology-based evaluation offers new insights into tractography algorithm performance. While probabilistic methods tend to enhance morphological consistency across cohorts, their broader spatial coverage may obscure subtle pathological differences. Balancing this trade-off is critical when selecting tractography approaches, especially in clinical applications where detecting fine-grained neuroanatomical changes is essential for disease diagnosis and monitoring progression.

## Acknowledgment

We thank the ENIGMA-Tractometry working group for providing access to data resources and analytic tools. Research reported in this publication was supported by NIH grants R01AG060610, RF1AG057892, R01MH134004.

